# From wag to affect: Tail kinematic signatures of canine emotional states in computer-controlled environments

**DOI:** 10.64898/2026.03.01.708848

**Authors:** Yuri Ouchi, Cara Glynn, Chiara Canori, Sarah Marshall-Pescini, Fumihide Tanaka, Friederike Range, Tiago Monteiro

## Abstract

From facial expressions to gestures, animals use multiple signal modalities to express emotions and communicate. In dogs, tail movements are conspicuous behaviours associated with emotional states, but this link remains debated. We investigated canine emotional states underlying tail wagging by systematically analysing differences in tail movements in a computer-controlled task encompassing two non-social *Conditions – Rewarded* (positive) *and Unrewarded* (negative), and two *Epochs* (pre-response and outcome anticipation). Using pose-tracking we found that 11 out of 23 dogs did not wag their tails in at least 75% of trials, suggesting that some dogs may inherently wag less or that tail wagging is primarily a social signal. Our results showed that dogs were more likely to wag during positive anticipation; whereas in the negative condition, despite tail amplitude being more prominent, increased speeds reflected arousal rather than valence. Further work should assess tail kinematics in social contexts to test and extend these findings.

## INTRODUCTION

Domestic dogs display visual, auditory, and olfactory cues in social interactions (Miklosi, 2014; Range and Marshall-Pescini, 2022) as well as in non-social contexts such as interactions with objects or food rewards (Flint et al., 2018; Prato-Previde et al., 2023). Visual signals such as facial expressions (Kaminski et al., 2017; Pedretti et al., 2022, 2024), behavioural displays (e.g., head turning, nose licking, yawning) (Mariti et al., 2017; Pedretti et al., 2023), and tail movements (Simpson, 1997; Quaranta, Siniscalchi and Vallortigara, 2007; Siniscalchi et al., 2013, 2018; Leonetti et al., 2024) are frequently used by humans to interpret dogs’ behaviour and underlying emotional states (Handelman, 2012; Serpell, 2017). In particular, tail wagging is regarded as one of the most prominent and widely recognised indicators when categorising dogs’ emotional behaviours (Tami and Gallagher, 2009; Leonetti et al., 2024) and has mostly been measured based on its occurrence (Travain et al., 2016; Pedretti et al., 2024) or duration (Flint et al., 2018; Prato-Previde et al., 2023).

Previous studies have shown that specific tail movements may correlate with putative emotional states in dogs. For example, loose wagging was assumed to indicate affiliative behaviours, while stiff tails were interpreted as a sign of anxiety (Fox and Bekoff, 1975). Additionally, tail height was suggested to play a critical role and be associated with emotional valence and/or social status, with high tail positions described as indicative of dominance and/or confidence, while low tail positions were associated with submissiveness and/or fear (Simpson, 1997; Serpell, 2017).

Furthermore, left-right asymmetries in tail wagging have emerged as yet another indicator of dogs’ emotional states. Quaranta and colleagues measured tail wagging angles and showed that dogs displayed right-biased tail wagging when facing humans, whereas left-biased tail wagging was observed in interactions with unfamiliar dominant dogs, suggesting that right-biases might reflect (positive) approach tendencies while left-biases might be associated with (negative) withdrawal tendencies related to brain lateralisation (Quaranta et al., 2007). Consistent with this study, Siniscalchi et al. (2013) found that dogs that observed left-biased wagging in another dog had increased heart rates and showed signs of anxiety. In addition, temporal shifts in tail-wagging asymmetry, from left to right, were observed over three days of brief interactions between laboratory beagles and an unfamiliar experimenter (Ren et al., 2022), suggesting that experience with positive events might induce rightward biases in tail wagging. Contrary to these results, Artelle and colleagues showed that dogs that observed a robot dog displaying a right-biased wagging were more likely to stop approaching it, while dogs were more likely to approach the robot when observing left-biased tail wagging, questioning the directionality of tail wagging asymmetry and its relation with brain lateralization (Artelle et al., 2011). Moreover, in an olfaction task, dogs displayed greater leftward tail wagging amplitude when in the target odour area, despite the expectation that successful target detection would be perceived as positive reinforcement and hence should have induced a right-biased tail wagging (Martvel and Pedretti et al., 2025). Overall, the existing literature is limited and illustrates that various features of tail movement – including changes in speed, height and/or lateralisation – may act as a means for signalling emotional or affective states, yet which specific states they index remains unclear. Building on this evidence, more fine-grained analyses of specific tail-movement parameters could help clarify if and how these components indeed relate to dogs’ emotional states - an endeavour that has been challenging to pursue in previous studies for various reasons (**Table 1**, Problems).

**Table 1.**
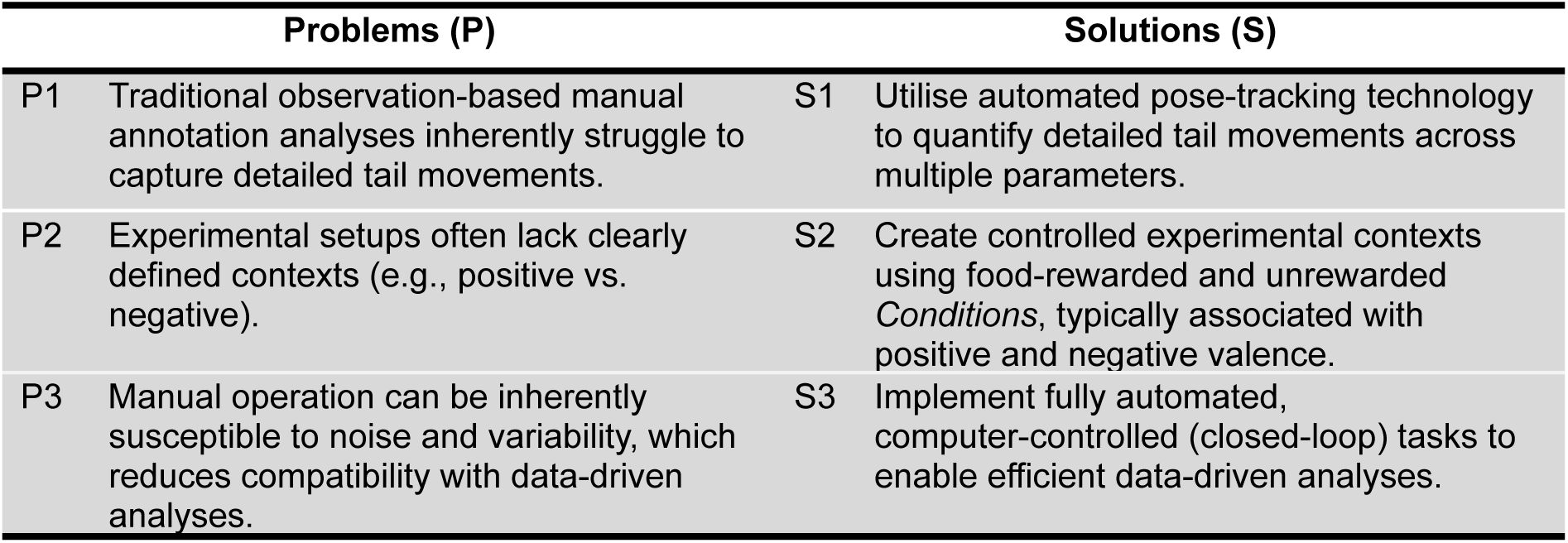
Summary of common problems identified in prior studies and corresponding solutions implemented in this study.

| Problems (P) |  | Solutions (S) |  |
| --- | --- | --- | --- |
| P1 | Traditional observation-based manual annotation analyses inherently struggle to capture detailed tail movements. | S1 | Utilise automated pose-tracking technology to quantify detailed tail movements across multiple parameters. |
| P2 | Experimental setups often lack clearly defined contexts (e.g., positive vs. negative). | S2 | Create controlled experimental contexts using food-rewarded and unrewarded *Conditions*, typically associated with positive and negative valence. |
| P3 | Manual operation can be inherently susceptible to noise and variability, which reduces compatibility with data-driven analyses. | S3 | Implement fully automated, computer-controlled (closed-loop) tasks to enable efficient data-driven analyses. |

Observation-based analyses lack the capacity to capture complex tail kinematics, which are composed of multiple parameters (**Table 1**, P1). This critical disadvantage can be solved by recent progress in machine learning-based markerless animal pose tracking (e.g., Mathis et al., 2018; Pereira et al., 2022) which has enabled non-invasive automatic quantification of animal behaviour (e.g., without the need for wearable devices such as accelerometers (Aich et al., 2019)), significantly reducing the time required for manual annotation (Quaranta et al., 2007) and minimising observer biases (**Table 1**, S1). Despite these advancements, few studies to date have applied tracking-based computational methods to canine behavioural and cognitive research and mainly focused on either detecting specific actions (e.g., Barnard et al., 2016) or emotions (e.g., Ferres, Schloesser and Gloor, 2022; Martvel and Riemer, 2025) using static images. The ones that have tracked body and tail movements from video (Martin et al., 2022, 2024; Ren et al., 2022; Völter et al., 2023a; Völter et al., 2023b; Martvel and Pedretti et al., 2025) were mostly unable to assess and interpret the impact of emotional valence on such movements due to the experimental environments lacking clearly parsable contexts (e.g., positive vs. negative – **Table 1**, P2).

Pedretti and colleagues (2024) investigated dogs’ facial expressions and other behavioural differences in a frustration-inducing contextual experiment where inaccessible food was displayed and then moved towards a partner (human or dog – *Social Condition*) or to an empty space (*Non-social Condition*), with dogs exhibiting clear differential facial patterns across *Conditions* and also increased their tail wagging frequency in the *Social Conditions* compared to the *Non-social* one. However, the experiment was not designed to track tail movements in detail, thus providing no information on whether tail wagging movement profiles (including lateral biases) changed depending on whether the stimulus was a human or an unfamiliar dog, or even across social and non-social conditions. This highlights the challenge of applying data-driven analyses that combine subtle movements with task events. The problem arises from the fact that manual task-deployment is inherently susceptible to increased noise and variability, which impact data-driven approaches that apply standardised treatments to the data (**Table 1**, P3).

Here we set out to decipher dogs’ emotional states underlying tail movements by investigating the impact of experimentally controlled contextual changes (i.e., positive and negative events) in dogs’ subtle tail movements by precisely quantifying multiple tail parameters while addressing limitations associated with traditional manual task deployment and control methods. We defined two *Conditions* where dogs were either food-rewarded or unrewarded, creating two contexts that were easily parsable to be combined with high speed video data (**Table 1**, S2). The experimental paradigm was adopted from Bremhorst and colleagues (2019) who observed different patterns of dogs’ facial expressions between positive-anticipation and frustration contexts based on whether or not food was delivered (see also Pedretti et al., 2022, 2024). Additionally, we employed markerless pose-tracking technology (**Table 1**, S1; Mathis *et al*., 2018) and a fully automated (i.e., closed-loop), computer-controlled task to facilitate data-driven analyses while reducing noise and variability during data acquisition (**Table 1**, S3; **Figure 1A, B**).

**Figure 1.**
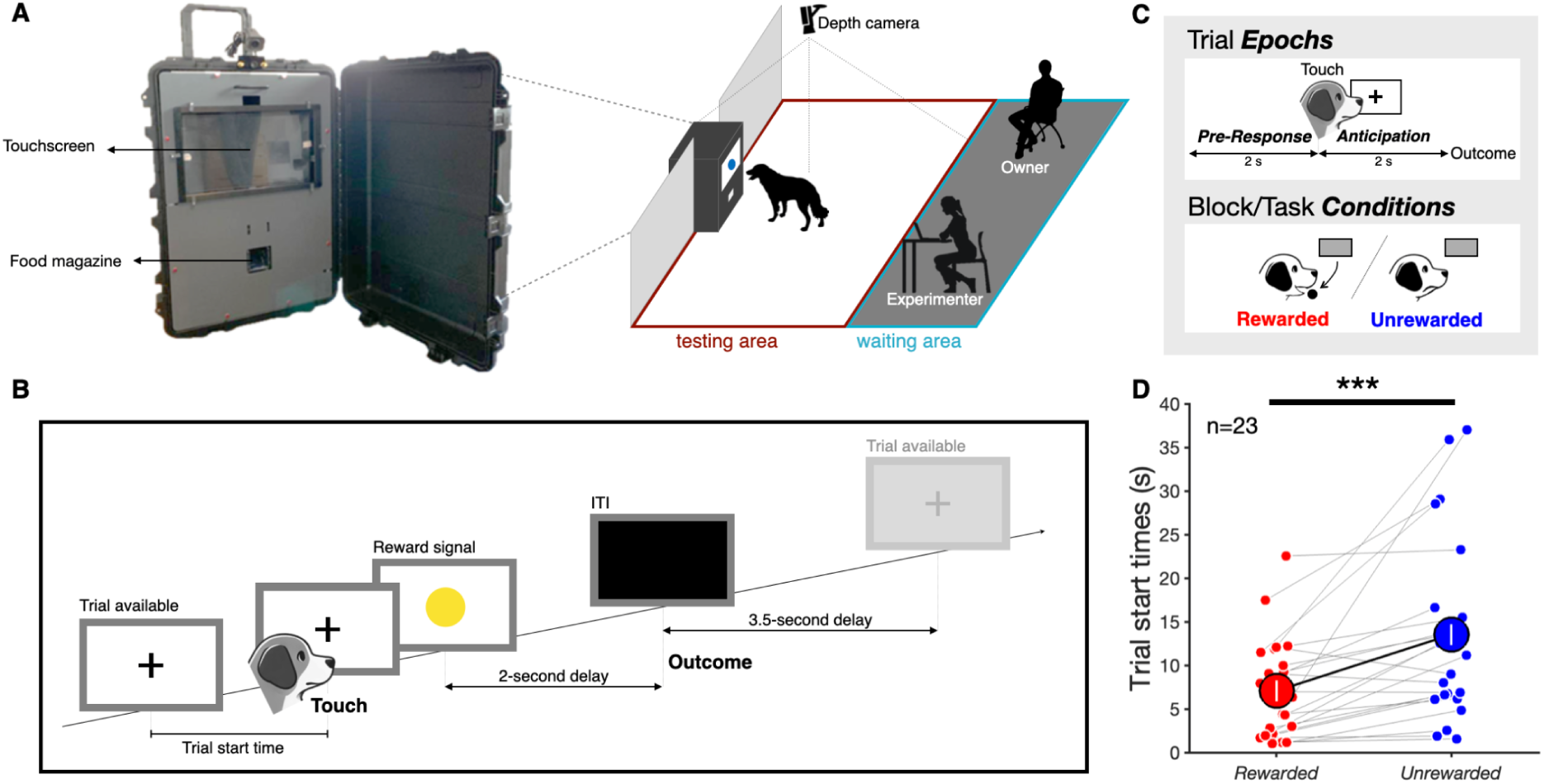
Experimental framework. **(A)** *Left:* Experimental setup. The touchscreen-based testing apparatus included an infrared touchscreen interface, a 17’’ computer screen and an automated food dispenser. Dry food pellets were dispensed automatically from the opening located in the bottom of the device (food magazine). *Right:* Experimental environment. The experimental room was split into two areas. The dog owner and experimenter were in the area close to the room’s entrance (*waiting area*). *The testing area* was in the middle of the room and constituted the space allocated to the testing apparatus and the subject dog, both completely in the field view of an overhanging depth camera. **(B)** Trial flow. There were five different events in a testing trial (*Trial available* – *Touch* – *Reward Signal* – *Outcome – Inter-Trial Interval (ITI*)). A touch to the screen during *Trial available* triggered the immediate presentation of the *Reward Signal* followed by either a reward (*Rewarded Condition*) or no reward (*Unrewarded Condition*) after a 2-second presentation period. The ITI lasted 3.5 seconds immediately following the outcome (reward or no reward) and preceding a new trial. **(C)** Top: for analyses, we defined 2-second *Epochs* before (*Pre-Response Epoch*) and after (*Anticipation Epoch*) a dog initiated any given trial (see *Experimental Procedure* for details). Bottom: the task followed a 6-trial block structure for *Rewarded* and *Unrewarded* conditions. **(D)** Trial start times for *Rewarded* (red) and *Unrewarded* (blue) block *Conditions*. Larger markers show the means across all subjects (±SEM; *n* = 23; \*\*\**p* < 0.001) and small markers (laterally displaced for visualisation purposes) indicate individual animal means. Grey lines connect the same individuals’ values.

Dogs were initially trained to start trials with a touch to the screen (i.e., trial start response), followed by the pairing of an arbitrary stimulus (i.e., a coloured circle presented on the computer screen) with a food reward (**Figure 1B**). In order to manipulate the experienced valence (positive vs. negative), in the *Rewarded Condition*, a food reward always followed the trained contingency, while in the *Unrewarded Condition*, this contingency was violated; despite the same stimulus being presented, no food reward was delivered (**Figure 1C**, bottom). Anticipation states are known to increase activity levels in animals (Boissy *et al*., 2007; McGowan *et al*., 2014), highlighting the need for comparing the tail movements across task epochs, in addition to task conditions. To this extent, our experimental contexts, positive and negative, comprised two distinct, non-overlapping, continuous task *Epochs* used for analyses (**Figure 1C**, top): aligned on a dog’s screen touch to initiate a given trial, the *Pre-Response Epoch* was defined as the 2-second interval before this response (but after the previous trial outcome was revealed), and the *Anticipation Epoch,* defined as the, 2-second interval following the same response (and before the current trial outcome was revealed).

Based on the 2 x 2 comparison (*Condition* * *Epoch*), we predicted that:

1. Dogs would exhibit differential tail movements across *Conditions*, reflecting the impact of being *Rewarded versus Unrewarded*.
2. Dogs would display more extreme tail movements during the *Anticipation Epoch* by increased tail parameter values compared to the *Pre-Response Epoch*.

To test these predictions, we used a block structure design, with *Conditions* alternating in blocks composed of 6 *Rewarded* or *Unrewarded* trials (adapted from Bremhorst et al., 2019).

This combination of parsable experimental contexts, technology-enhanced experimental operation and data analyses allowed for complex tail movements to be decomposed into wagging occurrence (a binary metric) and six (continuous) tail parameters (i.e. angle as a tail wagging asymmetry indicator, rhythmicity, amplitude, velocity, acceleration, and height) and aimed to reveal subtle associations between dogs’ tail movements and emotional valence.

## RESULTS

### Trial start times

We used trial start times to confirm whether the task’s contingencies and block structure functioned as intended by eliciting distinct responses between the *Rewarded* and *Unrewarded* block *Conditions*. Indeed, dogs (*n* = 23) took longer to initiate trials in the *Unrewarded Condition* than the *Rewarded Condition* (*estimate*±*SE* = 0.928±0.177, *t* = 5.26, *p* < 0.001, see **Figure 1D**, *Experimental Procedure - Trial Start Times* and Supplementary Figure 1 for more details).

### Tail movement parameters

We obtained seven body points (nose, head, withers, back, tail base, tail middle, and tail tip, Figure 2A) using DeepLabCut (Mathis et al., 2018). Prior to quantifying tail movement parameters, we examined the quality of the pose-tracking output. Five animals were excluded due to a high proportion of missing data caused by poor automated pose-tracking performance (Supplementary Table 1). Independent visual inspection of the corresponding videos confirmed this assessment revealing minimal to no tail movement during the experiment for these individuals (See *Data Selection* - *Tracking quality evaluation* for details). For the remaining 18 dogs, we estimated the tail angle relative to the dog’s body, by calculating the angle between two vectors: (1) the vector defined by the back and tail base points and (2) the vector defined by the tail base and the tail middle points. Negative to positive values indicate left to right positions (Figure 2A). Tail movements were first quantified by the change in tail angle, then tail wagging occurrence was detected in each trial by applying a Fourier transform to the time series of tail angle and subsequently decomposed and analysed based on five parameters (see Figure 2B and *Data Acquisition* - *Tail Kinematic Parameters* for further details): tail angle (negative-left to positive-right, a metric for tail wagging asymmetry), amplitude (in degrees, the difference between adjacent positive and negative peaks in tail angle wave), velocity (in degrees per second), acceleration (in degrees per second squared), and rhythmicity (based on the entropy of the normalised power spectrum of tail angle). Rhythmicity was adopted as a tail parameter since tail wagging has been recently described as a stereotyped, cyclical, and rhythmic behaviour (Leonetti et al., 2024), despite also giving rise to arrhythmic patterns (of different tail parameters) over short periods of time, which also characterised tail movements. In addition, tail height was defined as the height of the tail middle point relative to the floor and obtained independently of the time series of tail angle as another tail parameter (see *Data Aquisition - Section Depth Data Alignment and Processing* for details).

**Figure 2.**
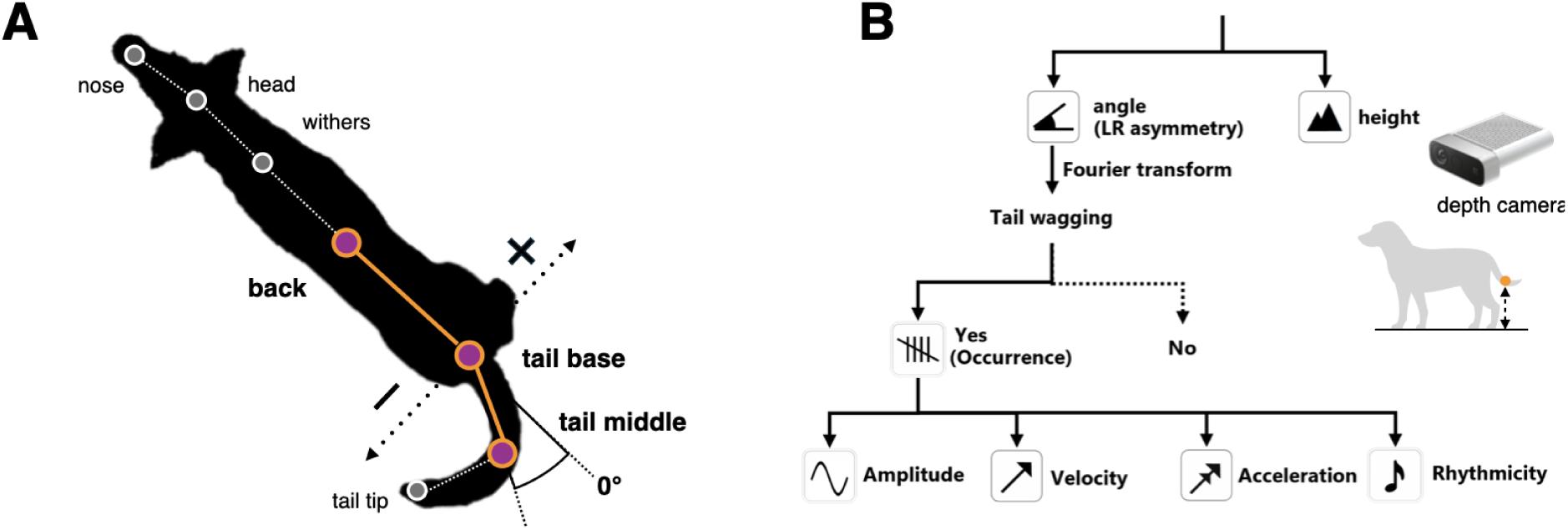
Tail tracking and movement parameters. **(A)** Tracked body nodes and tail angle. We obtained seven body nodes using DeepLabCut. As a proxy for tail angle relative to the dog’s body, we calculated the angle between two vectors: (1) the vector defined by *back* and *tail base* points and (2) the vector defined by the *tail base* and the *tail middle* points. Negative to positive values indicate left to right positions. **(B)** Tail movements were first abstracted by the change in tail angle relative to a dog’s body axis (see *Tail Kinematic Parameters* section for details). Tail wagging was decomposed and analysed based on six parameters: tail angle (negative-left to positive-right), amplitude (in degrees, the difference between adjacent positive and negative peaks in tail angle wave), velocity (in degrees per second), acceleration (based on velocity), and rhythmicity (based on the entropy of the normalised power spectrum of tail angle). Tail height data (relative to the ground) was obtained directly from a depth camera and aligned to the tracking data (see *Data Acquisition - Depth Data Alignment and Processing* for details).

#### Wagging occurrence

Among the 18 dogs with sufficient tracking data, we found no significant effect of *Conditions* or task *Epochs* on the probability of tail wagging occurrence (full-null model comparison *χ²*(3) = 3.94, *p* = 0.268; Supplementary Figure 2). However, 6 dogs wagged their tails in fewer than 25% of trials across *Conditions* and task *Epochs* (Figure 3A) and were thus excluded from further analyses. The remaining 12 dogs formed *the wagging group (*minimum wagging occurrence ratio > 0.25; threshold defined by visual inspection of the data, see Figure 3A). In this *wagging group* (*n =* 12), we found an interaction between *Condition* and task *Epoch* in the probability of wagging occurrence (full-null model comparison *χ²*(3) = 7.97, *p* < 0.05; interaction *Condition-Epoch*: AIC = 2820, LRT = 4.80, *p* < 0.05; Figure 3B; see also Supplementary Figure 3 for within *Condition* comparisons).

**Figure 3.**
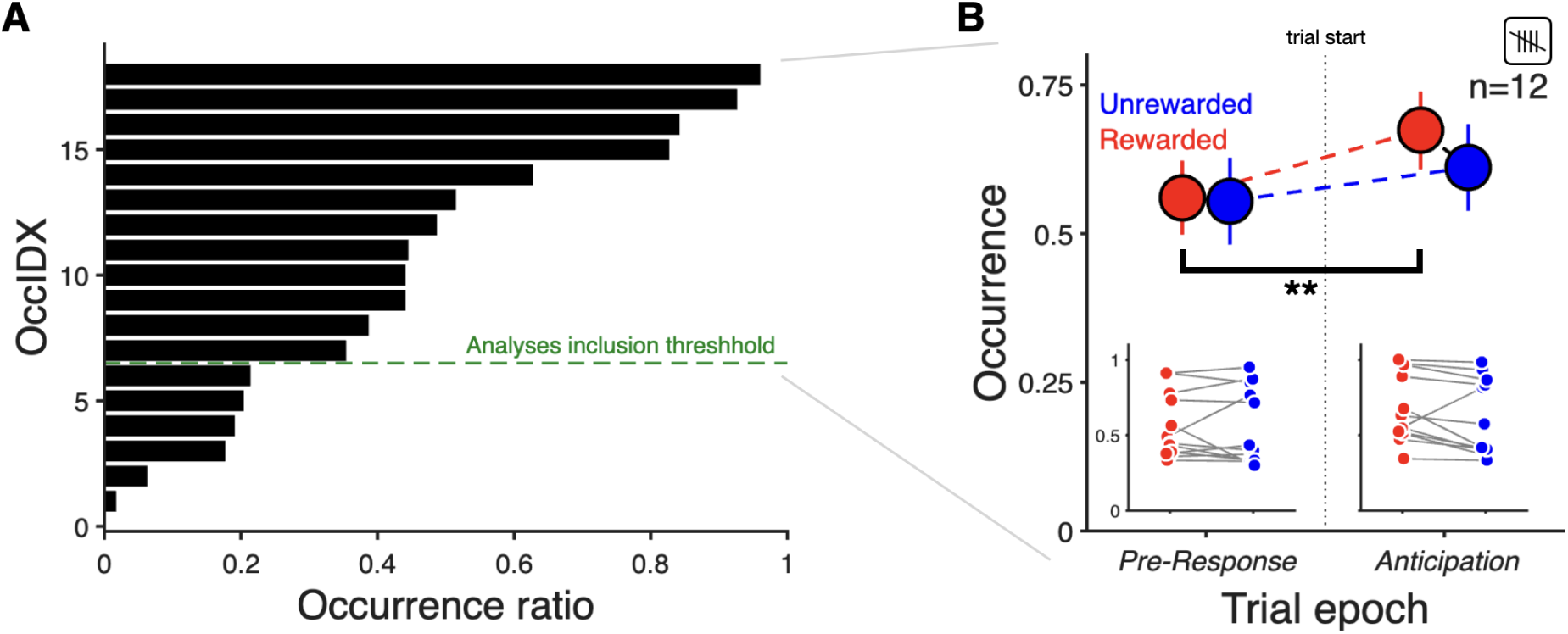
Tail-wagging occurrence metrics. **(A)** Overall ranking for tail-wagging occurrence. The histogram shows tail-wagging occurrence ratios across *Epochs*, calculated by dividing the number of trials in which tail wagging occurred by the total number of valid trials. The green dashed line separates the 12 dogs (above) that were included in further analyses and the ones (below) that were excluded due to reduced tail movements (see Supplementary Table 1 and Supplementary Figure 2 for more details). **(B)** Tail-wagging occurrence ratio across task *Conditions* and *Epochs* for dogs above the minimal occurrence threshold (*wagging group*). Red and blue markers show tail-wagging occurrence, aligned to a trial start response (vertical dotted line) from *Rewarded* and *Unrewarded Conditions* across *Pre-Response* and *Anticipation Epochs*, respectively. Larger markers indicate the means across all subjects (±SEM; *n =* 12; ** *p* < 0.01). Insets show individual mean values (laterally displaced for visualisation purposes) and grey lines connect the same individuals’ values across block *Conditions* and trial *Epochs*. See also Supplementary Figure 2 for the same analysis applied to all dogs irrespective of their occurrence index, and Supplementary Figure 3 for within *Condition* comparisons).

To further investigate this interaction, we estimated marginal means (EMMs) with a Bonferroni correction for multiple comparisons (1: the effect of *Epoch* within each level of *Condition*, 2: the effect of *Condition* within each level of *Epoch*). This post-hoc analysis indicated that dogs wagged their tails more often during the *Anticipation Epoch* than the *Pre-Response Epoch* in the *Rewarded Condition* (Odds Ratio = 2.44, *p* < 0.01).

#### Angle

We found no effects of *Condition* and *Epoch* on tail angles (i.e., no left-right asymmetries; *n =* 12; full-null model comparison *χ²*(3) = 4.18, *p* = 0.243; Figure 4A; see also Supplementary Figure 3 for within *Condition* comparisons).

**Figure 4.**
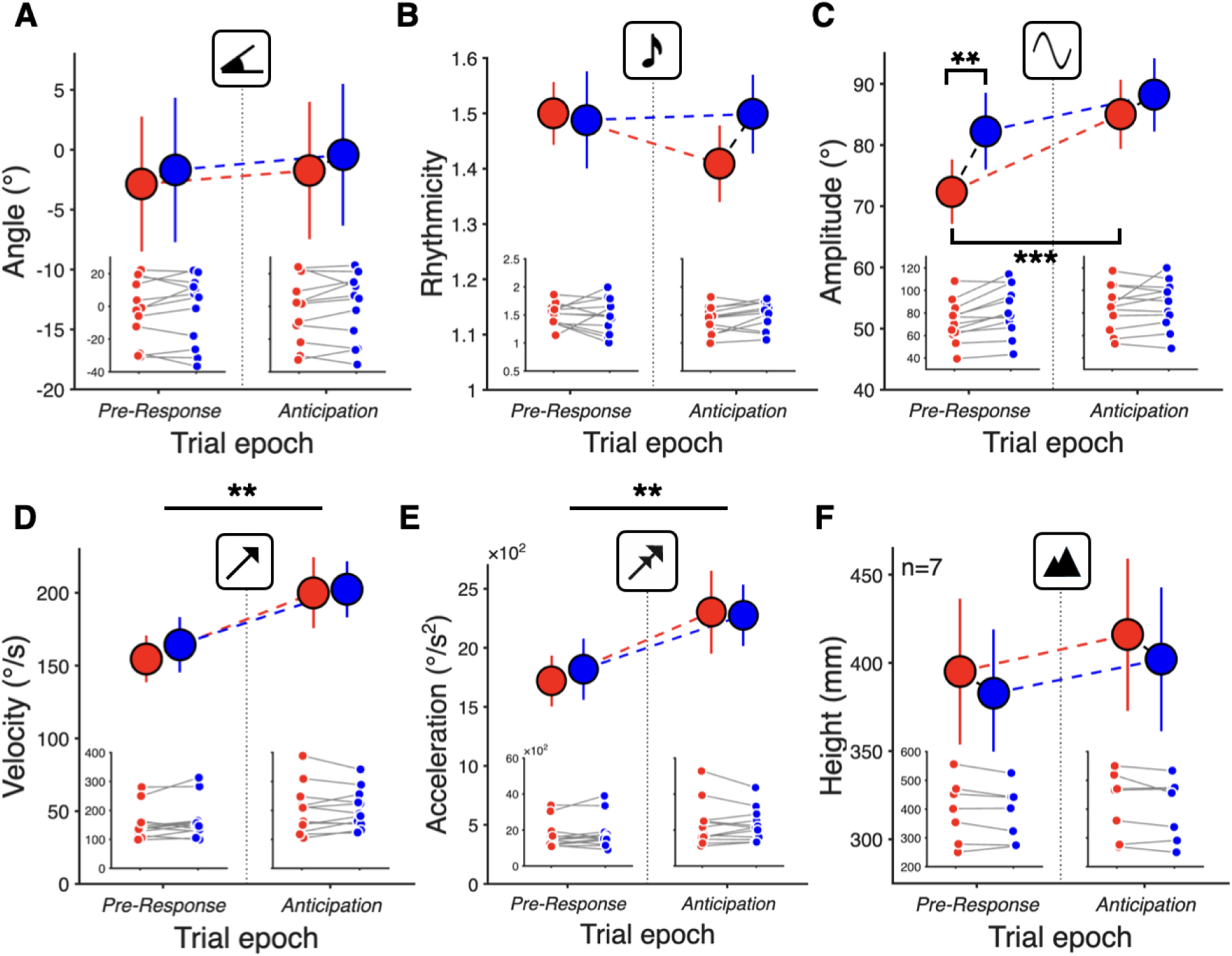
Continuous tail movement parameters’ results. A to. **F.** Red and blue markers show tail parameter data, aligned to a trial start response (vertical dotted line) from *Rewarded* and *Unrewarded Conditions* across *Pre-Response* and *Anticipation Epochs*, respectively. Y-axes labels (and icons, see Figure 2B) indicate the corresponding tail parameter with larger markers indicating the means across all subjects (±SEM; *n* = 12 for all parameters except Height (*n* = 7); ** *p* < 0.01, *** *p* < 0.001). Insets show individual mean values (laterally displaced for visualisation purposes) and grey lines connect the same individuals’ values across block *Conditions* for each trial *Epoch*.

#### Wagging rhythmicity

The full-null model comparison was not statistically significant (*χ²*(3) = 4.66, *p* = 0.199; Figure 4B; see also Supplementary Figure 3 for within *Condition* comparisons).

#### Wagging amplitude

Following a full-null model comparison (*χ²*(3) = 21.1, *p* < 0.001), we found a significant interaction between *Condition* and task *Epoch* in tail wagging amplitude (*estimate±SE* = 0.159±0.043, *t*(23.2) = 3.70, *p* < 0.001; Figure 4C; see also Supplementary Figure 3 for within *Condition* comparisons). A post-hoc analysis using EMMs revealed that wagging amplitude was significantly higher during the *Anticipation* than the *Pre-Response Epoch* in the *Rewarded Condition* (*estimate±SE* = 0.224±0.036, *t.ratio*(10.3) = 6.23, *p* < 0.001). In the *Pre-Response Epoch,* wagging amplitude was significantly higher in the *Unrewarded* than the *Rewarded Condition* (*estimate±SE* = −0.169±0.047, *t.ratio*(10.2) = −3.59, *p* < 0.01).

#### Wagging velocity & acceleration

We found that velocity and acceleration were higher in the *Anticipation* relative to the *Pre-Response Epoch* (Velocity: full-null model comparison *χ²*(3) = 15.7, *p* < 0.001; *estimate±SE* = −0.260±0.067, *t*(11.6) = −3.90, *p* < 0.01; Acceleration: full-null model comparison *χ²*(3) = 14.4, *p* < 0.01; *estimate±SE* = −0.238±0.059, *t*(11.5) = −4.06, *p* < 0.01), whereas the main effect of *Condition* was not significant for neither velocity (*estimate±SE* = 0.005±0.05, *t*(12.5) = 0.105, *p* = 0.918) nor acceleration (*estimate±SE* = 0.013±0.053, *t*(10.7) = 0.238, *p* = 0.817) (Figure 4D**,E**; see also Supplementary Figure 3 for within *Condition* comparisons).

#### Height

Five dogs were excluded from the dataset due to unreliable depth data (see *Data Selection* - *Curated datasets* for details). Despite a non-significant full-null model comparison (*n* = 7; *χ²*(3) = 2.98, *p* = 0.394), tail heights were consistently higher in the *Rewarded Condition* than the *Unrewarded Condition*, and also during the *Anticipation* rather than the *Pre-Response Epoch* (Figure 4F; see also Supplementary Figure 3 for within *Condition* comparisons).

## DISCUSSION

In this study, we examined the emotional basis of tail wagging in dogs using a self-paced, computer-controlled task, that compared *Food-Rewarded* and *Unrewarded Conditions*, assuming these differed in emotional valence (i.e., positive vs. negative). Firstly, the effectiveness of the experimental paradigm was confirmed with trial start times being shorter, on average, in the *Rewarded* than in the *Unrewarded Condition* (Figure 1D), a result consistent with dogs’ having learned the basic task contingencies. Secondly, we found that only 12 dogs wagged their tail in more than 25% of the trials across our *Conditions* (Figure 3A). In these 12 dogs, we observed more pronounced tail movements during the *Anticipation Epoch* reflected in increased tail wagging velocity and acceleration (Figure 4D**,E**; **Table 2**). Finally, we found an interaction between task *Condition* and *Epoch* in both wagging occurrence and amplitude (Figures 3B and **4C**, respectively; see also **Table 2**).

**Table 2.** Summary of Tail parameters’ results.

| ***Tail Parameters*** | ***n*** | ***Condition*** | ***Phase*** | ***Sig.*** |
| --- | --- | --- | --- | --- |
| Wagging Occurrence | 12 | *Rewarded* | *Anticipation > Pre-Response* | ** |
| Velocity & Acceleration | 12 | *Rewarded & Unrewarded* | *Anticipation > Pre-Response* | ** |
| Amplitude | 12 | *Unrewarded > Rewarded* | *Pre-Response* | ** |
|  |  | *Rewarded* | *Anticipation > Pre-Response* | *** |
| Angle | 12 | *n.s.* |  |  |
| Rhythmicity | 12 | *n.s.* |  |  |
| Height | 7 | *n.s.* |  |  |

note: \*\* $p < 0.01$ , \*\*\* $p < 0.001$

Out of 23 dog participants, approximately half of the dogs (*n =* 11) did not wag their tails in at least 75% of all trials, suggesting that these dogs may either inherently exhibit low levels of tail wagging regardless of the situation, or tail wagging may function primarily as a social communicative signal making our non-social experimental context not sufficiently relevant for eliciting detectable tail wagging movements.

Among the 12 reliable waggers, tail wagging occurred more often in the *Anticipation* than the *Pre-Response Epoch* within the *Rewarded Condition*, with no corresponding *Anticipation–Pre-Response* differences in the *Unrewarded Condition* and no overall main effect of *Condition* (Figure 3B; **Table 2**). Moreover, as we expected (*Prediction 2*), dogs’ wagging speed was higher during the *Anticipation* than during the *Pre-Response Epoch* (Figure 4D**,E**; **Table 2**). Once again, neither wagging velocity nor acceleration differed between *Conditions.* Taken together, these results suggest that dogs might be more likely to wag their tails in positive anticipation situations, but, when tail wagging is present in frustration situations, its velocity and acceleration indicate arousal rather than emotional valence. Thus, the presence of tail wagging could serve as a marker for positive arousal regardless of the presence of social partners (see also (McGowan *et al*., 2014; Travain *et al*., 2016)), while tail speed metrics might provide a more general arousal measure, irrespective of the valence of the corresponding situation.

Unexpectedly, wagging amplitude was higher during the *Pre-Response Epoch* in the *Unrewarded* than the *Rewarded Condition* and it was higher in the *Anticipation Epoch* than *the Pre-Response* within the *Rewarded Condition* (Figure 4C; **Table 2**). Together with the significant differences in wagging speeds (i.e., velocity and acceleration) between *Pre-Response* and *Anticipation Epochs,* these results suggest that amplitude may track emotional valence rather than arousal: in the negative context (*Unrewarded*), wagging amplitude remained elevated and did not mirror arousal-related speed changes. Specifically, higher amplitude wagging at moderate speed may index negative affective states, such as frustration.

With regards to tail height, despite the lack of statistical differences (Figure 4F; **Table 2**) likely due to the small sample size, we observed an increase in tail height from *Pre-Response* to *Anticipation Epochs* in the *Unrewarded Condition* in 6 out of 7 animals (Supplementary Figure 3). This trend suggests that tail height might be more strongly affected in negative situations, especially when the outcome is not fully predictable.

Interestingly, we found no effect of our two *Conditions* on tail wagging asymmetry (estimated from tail angle; **Table 2**) unlike what was reported in earlier studies (Quaranta et al., 2007; Siniscalchi *et al*., 2013; Ren *et al*., 2022; Martvel and Pedretti *et al*., 2025), which might be due to our adoption of a non-social testing context. A recent study also reported the absence of tail-wagging asymmetry in dogs tested in a non-social positive anticipation context associated with food or toys (Simon et al., 2024). Together these studies suggest that non-social contexts can attenuate dogs’ emotional dynamics and the associated changes in tail movements, with tail-wagging asymmetries occurring mainly in social contexts. Nevertheless, Martvel and Pedretti et al. (2025) reported tail asymmetries in a detection task; however, whether this paradigm is best classified as non-social or social remains unclear, as detection tasks typically involve a handler cue to start and an expected play interaction with the handler upon locating the target.

Overall, despite our across-condition trial start time differences (Figure 1D), it is possible that, against our initial prediction (*Prediction 1*), our task *Conditions* were not salient enough to induce differential tail movements when compared to differences between social and non-social situations (**Table 2**). In order to increase the contrast between *Rewarded* and *Unrewarded Conditions*, future studies could train dogs with two distinct stimuli, one for *Rewarded,* the other for *Unrewarded* outcomes respectively, allowing for trial randomisation (instead of predictable blocks) and a stable 50% reward probability throughout experimental sessions. Furthermore, since *Condition* blocks and associated trial *Epochs* were both relatively brief (i.e., *Condition* block was switched every 6 trials and each *Epoch* lasted 2 seconds; Figure 1C), these durations may not have been sufficiently long lasting to elicit detectable tail movement changes. Notably, studies that reported tail wagging asymmetries used longer trial durations (e.g., 60-s for stimulus presentation in Quaranta et al. (2007) and 5-min social interactions per day in Ren et al (2022)), suggesting that tail wagging asymmetries might only be observed in situations marked by extended temporal contingencies. Future studies should explore this matter further.

Moreover, including a neutral *Condition* may also help clarify the direction of change in each parameter relative to a neutral baseline. Although we focused on 2-second *Epochs* before and after responses (removing the food consumption period), additional *Condition*-specific behaviours may have occurred during and after the inter-trial-interval). Therefore, in order to detect differences, adopting unsupervised or semi-supervised clustering methods commonly used in neuroscience (e.g., Luxem *et al*., 2023; Tillmann *et al*., 2024; Weinreb *et al*., 2024) could be highly effective analyses alternatives due to their compatibility with pose-tracking technologies and behavioural classification capabilities. Additionally, as part of our selection of pose-tracking points, we used the tail middle as an anchor to characterize tail movements, as our data showed that it was a more robust landmark than the tail tip (i.e., with a lower percentage of missing values; see Supplementary Table 1). However, it is possible that other changes in tail parameters (beyond what we measured) remained undetected. Future studies may benefit from using more points along the tail to describe and model its movements in order to facilitate a more detailed and comprehensive characterisation of the resulting movement kinematics.

While we selected a ToF-based depth camera to capture 3D tail movements, this device proved to be unreliable for thin or black coloured tails. Notwithstanding, we confirmed that a single depth camera is sufficient for the acquisition of 3D information of different body points from dogs with different morphological characteristics (as long as they are not black or their tails too thin), overcoming the need to use multiple cameras and more complex triangulation and calibration procedures. While multiple cameras or wearable accelerometers can provide more accurate and stable data, a single depth camera, which is both easily deployable and non-invasive, enables testing across diverse environments and experimental contexts, including dog breeders and shelters, and could facilitate the monitoring and improvement of dog welfare.

Unlike conventional observation-based analyses that are inherently discrete, apply single standards to entire populations (e.g., Ruge *et al*., 2018), and thus potentially overlook variations within specific ranges for each parameter of interest, our approach highlighted substantial inter-individual variability in baseline measurements (see insets in Figure 3B and **4A-F**, and Supplementary Figures 2 and 3). For example, when defining tail height, categorising height above tail base as high and below it as low, may fail to account for changes occurring within the higher range. In contrast, the data-driven approach used in this study enabled the acquisition of continuous values at the individual level, which can allow for investigating variations across breeds and individual phenotypes, which has often been challenging in studies involving domestic dogs. The combined effectiveness of the controlled experimental setup and data-driven analyses should motivate further research, using a similar framework to investigate the relationship between dogs’ tail movements and putative emotional states in response to different contexts.

All in all, our study revealed that tail movements at least in non-social situations seem to mainly reflect arousal rather than emotional valence with the exception of amplitude being more prominent in negative contexts. Our computer-controlled experimental environment and pose-tracking-based data-driven analyses demonstrated its potential use in future studies aimed to investigate tail kinematic signatures of dogs’ putative emotional states in a time-efficient and unbiased fashion. Given our other findings of absence of 1) tail movements in half of our experimental subjects and 2) tail wagging asymmetries in the other half, future studies should combine a social aspect with our technology-enhanced experimental approach, aiming for a more detailed understanding of canine behaviour.

## RESOURCE AVAILABILITY

### Materials availability

- This study did not generate new unique reagents.

### Data and code availability

- Data reported in this paper will be shared by the corresponding contacts upon request.
- All original code will be deposited at repository and will be publicly available as of the date of publication.
- Any additional information required to reanalyze the data reported in this paper is available from the corresponding contacts upon request.

## ACKNOWLEDGMENTS

We would like to thank Giulia Pedretti, Hugo Marques, Svenja Capitain and all members of the Domestication Lab, past and present, for all the discussions and feedback. We would like to thank Karin Bayer and the Clever Dog Lab team for logistical support, Remco Folkertsma for statistical advice, the Champalimaud Hardware and Software platform (https://www.fchampalimaud.org/platforms/hardware-and-software-platform) and the Institute of Science and Technology Austria (ISTA) Miba Machine Shop (https://ista.ac.at/en/research/scientific-service-units/machine-shop/) from for help and support in building our experimental apparatus. We thank all dog owners for their availability to volunteer their dogs to participate in our experiment. Y.O. was funded by Ernst Mach Grant (weltweit), the Tobitate! Study Abroad Initiative and From Saitama to the World. F.R. and T.M. were supported by the Austrian Science Fund (FWF) Grant DOI 10.55776/P37052. C.C. was funded by a doctoral grant from the University of Parma (cycle XXXVIII). The study was also supported by a Research Grant from the Association for the Study of Animal Behaviour (ASAB) to T.M.

## AUTHOR CONTRIBUTIONS

Conceptualisation, Y.O. and T.M.; Methodology, Y.O., C.G., F.R. and T.M.; Investigation, Y.O., C.G., C.C. and T.M.; Writing—original draft, Y.O., F.R. and T.M.; Writing—review & editing, Y.O., C.G., C.C., F.T., F.R. and T.M.; Funding acquisition, Y.O., F.R. and T.M.; Resources, F.T., F.R. and T.M.; Supervision, F.T., F.R. and T.M.

## DECLARATION OF GENERATIVE AI AND AI-ASSISTED TECHNOLOGIES

We have not used AI-assisted technologies in creating this article.

## SUPPLEMENTARY INFORMATION

**Supplementary Table 1.** Missing value rates (%) within two task *Epochs* for each tracked body point after data processing. The white coloured records indicate the excluded dogs from the tail parameter analysis due to high missing value rates of tail middle.

| Dog ID | Tail tip | Tail middle | Tail base | Back | Withers | Head | Nose |
| --- | --- | --- | --- | --- | --- | --- | --- |
| 1 | 18.7 | 16.3 | 0.9 | 0.8 | 0.7 | 1.1 | 42.1 |
| 2 | 93.8 | 96.4 | 0.7 | 0.7 | 0.7 | 0.7 | 37.4 |
| 3 | 0.5 | 0.7 | 0.6 | 0.5 | 0.4 | 0.3 | 47.5 |
| 4 | 69.0 | 80.4 | 1.4 | 1.0 | 1.0 | 0.9 | 12.6 |
| 5 | 26.4 | 0.5 | 0.4 | 0.3 | 0.3 | 0.1 | 19.7 |
| 6 | 5.6 | 3.6 | 0.0 | 0.0 | 0.0 | 0.0 | 22.0 |
| 7 | 4.6 | 4.7 | 0.9 | 0.9 | 0.8 | 0.8 | 7.0 |
| 8 | 4.8 | 5.9 | 7.8 | 11.5 | 11.4 | 11.4 | 18.7 |
| 9 | 77.5 | 69.2 | 0.4 | 0.1 | 0.0 | 0.0 | 15.9 |
| 10 | 33.2 | 18.4 | 1.7 | 1.7 | 1.6 | 1.6 | 69.2 |
| 11 | 20.0 | 19.9 | 19.1 | 18.7 | 18.6 | 18.5 | 35.4 |
| 12 | 4.2 | 4.7 | 0.6 | 0.2 | 0.1 | 0.0 | 12.6 |
| 13 | 36.5 | 69.4 | 1.6 | 1.7 | 1.7 | 1.7 | 11.3 |
| 14 | 6.6 | 5.9 | 6.5 | 5.0 | 4.2 | 3.5 | 18.4 |
| 15 | 1.3 | 1.4 | 0.7 | 0.3 | 0.3 | 0.1 | 47.3 |
| 16 | 17.3 | 17.7 | 20.6 | 20.5 | 20.9 | 20.5 | 41.1 |
| 17 | 2.8 | 1.0 | 1.0 | 0.9 | 0.8 | 0.7 | 38.7 |
| 18 | 57.6 | 61.3 | 24.2 | 24.3 | 23.9 | 23.8 | 35.9 |
| 19 | 57.3 | 29.6 | 0.5 | 0.4 | 0.4 | 0.3 | 12.5 |
| 20 | 22.3 | 22.0 | 19.2 | 18.7 | 18.4 | 18.2 | 53.9 |
| 21 | 0.8 | 1.0 | 1.0 | 0.9 | 0.9 | 0.7 | 25.2 |
| 22 | 16.1 | 3.2 | 2.3 | 2.2 | 2.2 | 2.0 | 26.9 |
| 23 | 0.8 | 0.7 | 0.6 | 0.5 | 0.5 | 0.6 | 4.3 |

**Supplementary Figure 1.**
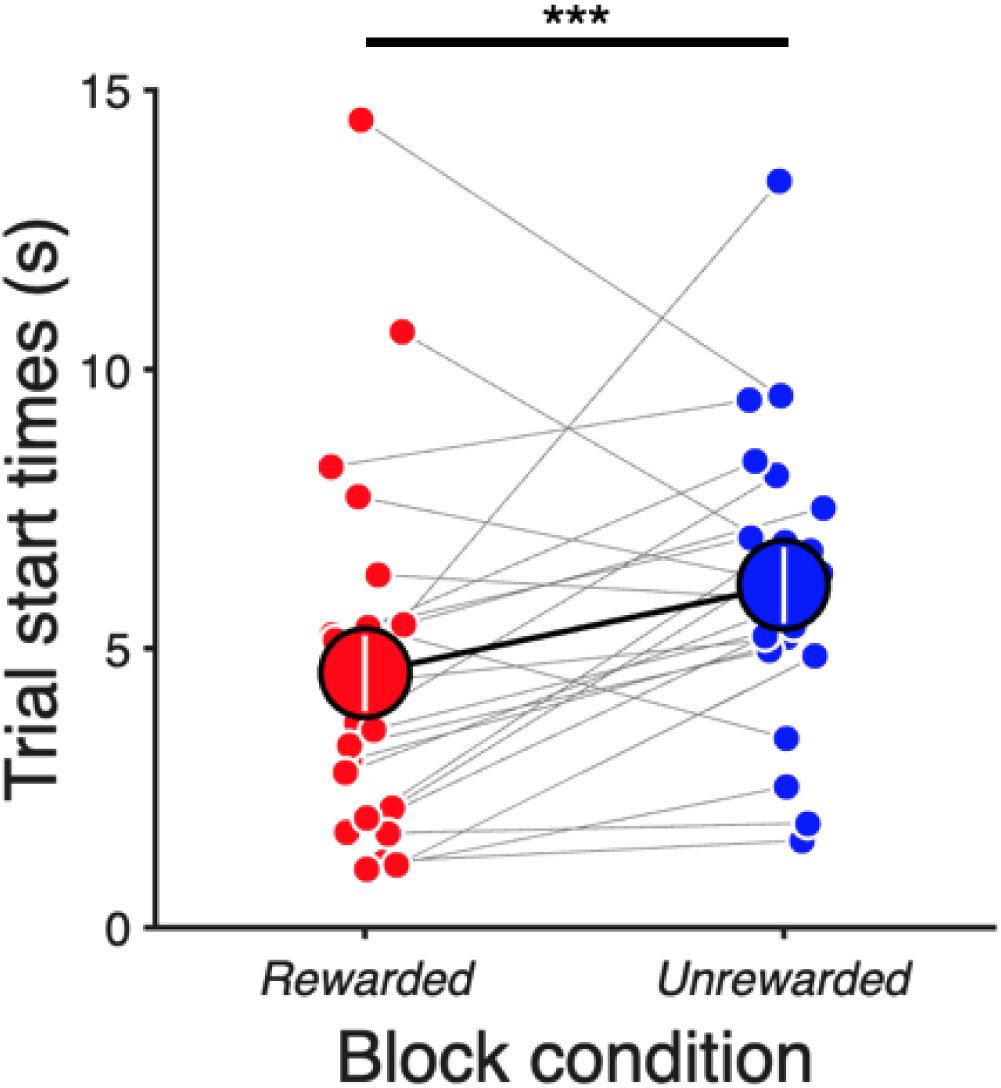
Trial start times excluding correction trials. Larger markers show the means across all subjects (±SEM; *n =* 23) and small markers indicate individual animal means (laterally displaced for visualisation purposes) for *Rewarded* (red) and *Unrewarded* (blue) block *Conditions*. Grey lines connect the same individuals’ values. (estimate±*SE* = 0.620±0.155, *t =* 4.00, *p <* 0.001).

**Supplementary Figure 2.**
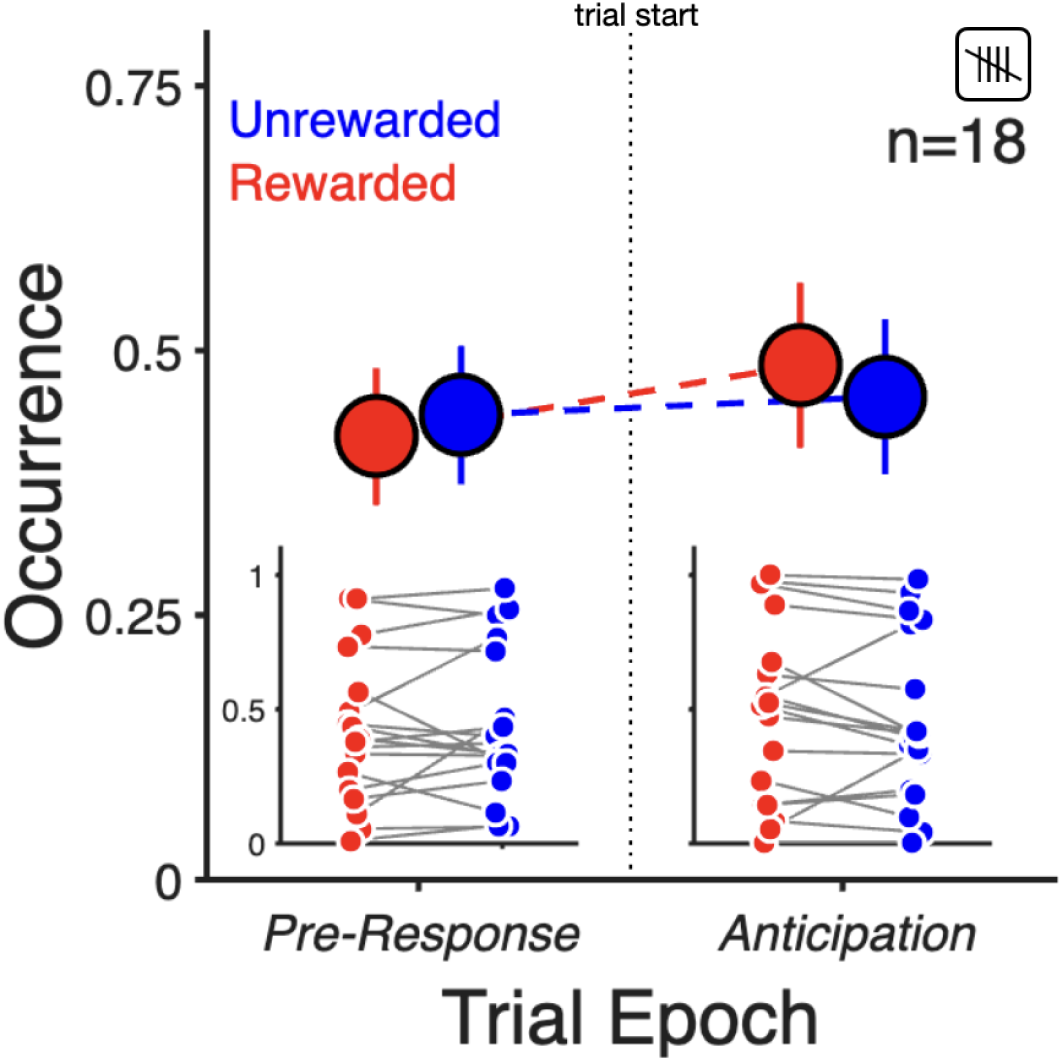
Tail-wagging occurrence ratio across task *Conditions* and *Epochs* (*n =* 18). Red and blue markers show tail-wagging occurrence data, aligned to a trial start response (vertical dotted line) from *Rewarded and Unrewarded Conditions* across *Pre-Response* and *Anticipation Epochs,* respectively; larger markers indicate the means across all subjects (±SEM; *n =* 18; *n.s.*). Insets show individual mean values (laterally displaced for visualisation purposes) and grey lines connect the same individuals’ values across block *Conditions* and trial *Epochs*. Same data presented in Figure 3B but now including the dogs that did not meet the minimal 0.25 occurrence threshold.

**Supplementary Figure 3.**
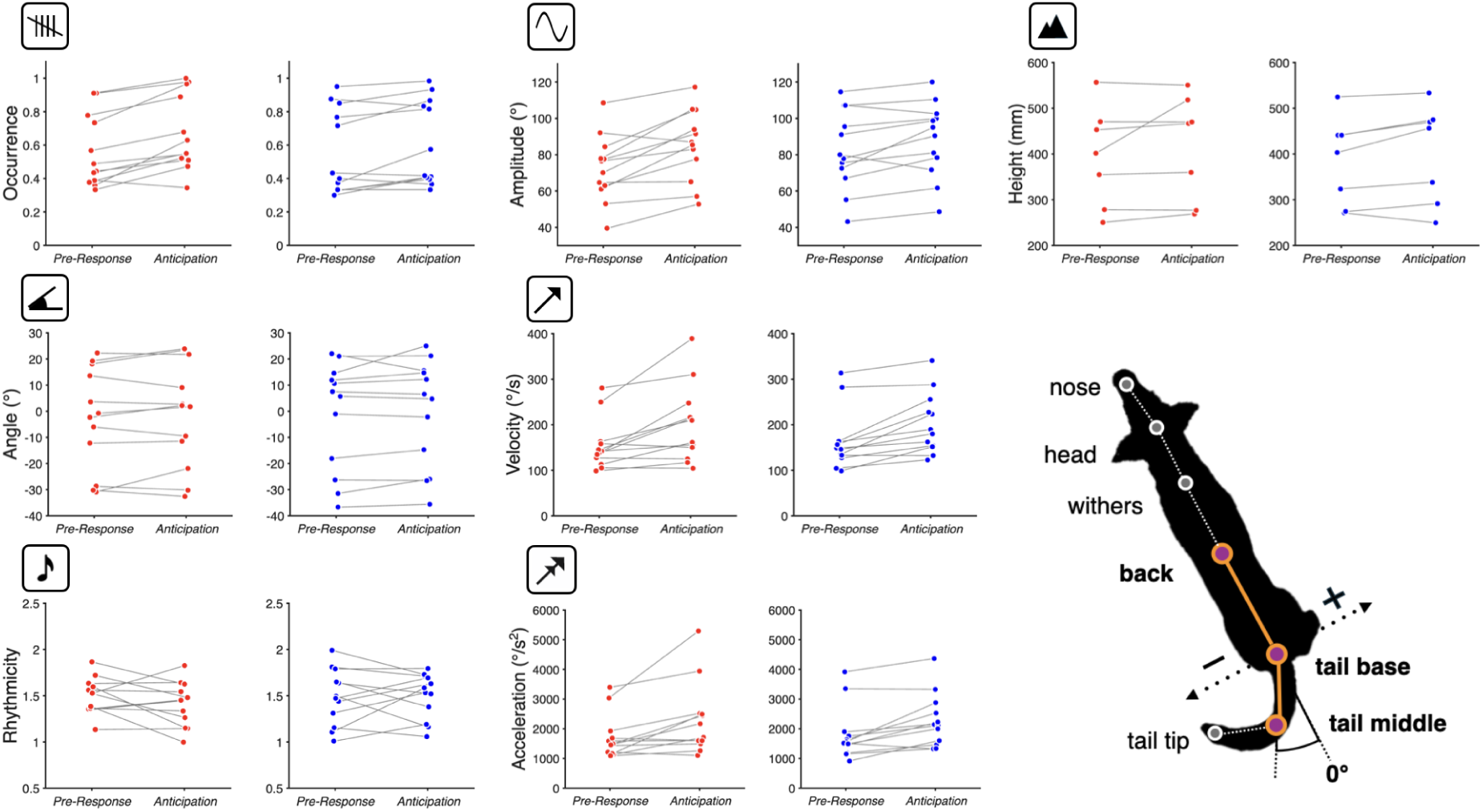
Tail tracking and movement parameters across task *Epochs* in each *Condition*. Same data presented in Figure 3B (wagging occurrence) and Figure 4 (remaining metrics), but highlighting within *Condition* comparisons across task *Epochs* (*n =* 12, except tail height (*n =* 7)). Y-axes (and icons) indicate the corresponding tail parameter. Markers depict individual mean values (laterally displaced for visualisation purposes) and grey lines connect the same individuals’ values across trial *Epochs* for each task *Condition* (*Rewarded*, red; *Unrewarded*, blue).

**Supplementary Video. Example tracking.** Example videos of different participants with overlaid DeepLabCut tracking. Subject IDs are 17, 3, 16, 22, 21, 1, 19, 15, 23 starting from the top left video.

## EXPERIMENTAL MODEL AND STUDY PARTICIPANT DETAILS

30 dogs were recruited from the dog owner database of the Clever Dog Lab, Austria, and social media. The subjects met the following criteria: having a long, straight tail and a nose height of at least 40 cm from the ground (i.e., when the head is in a relaxed position) to facilitate interacting with the touchscreen on the testing apparatus (**Figure 1A**, left). 7 out of 30 dogs were excluded due to low food motivation and/or not completing training. We used 23 pet dogs (12 males and 11 females; mean age: 6 years (or 75 months), range: 20–143 months) for the initial (trial start time) analysis. Among these, 12 dogs were classified as shepherd types, 3 as hunting types, 3 as guarding types, 1 as a retrieving and flushing type, and 4 as mixed breeds whose types could not be determined. More details of the individual dogs can be found in **Individual dog information**, below.

### Individual dog information

Participants are ordered by decreasing wagging occurrence ratio (Occ IDX). Dog ID remains the same as used for statistical analyses.

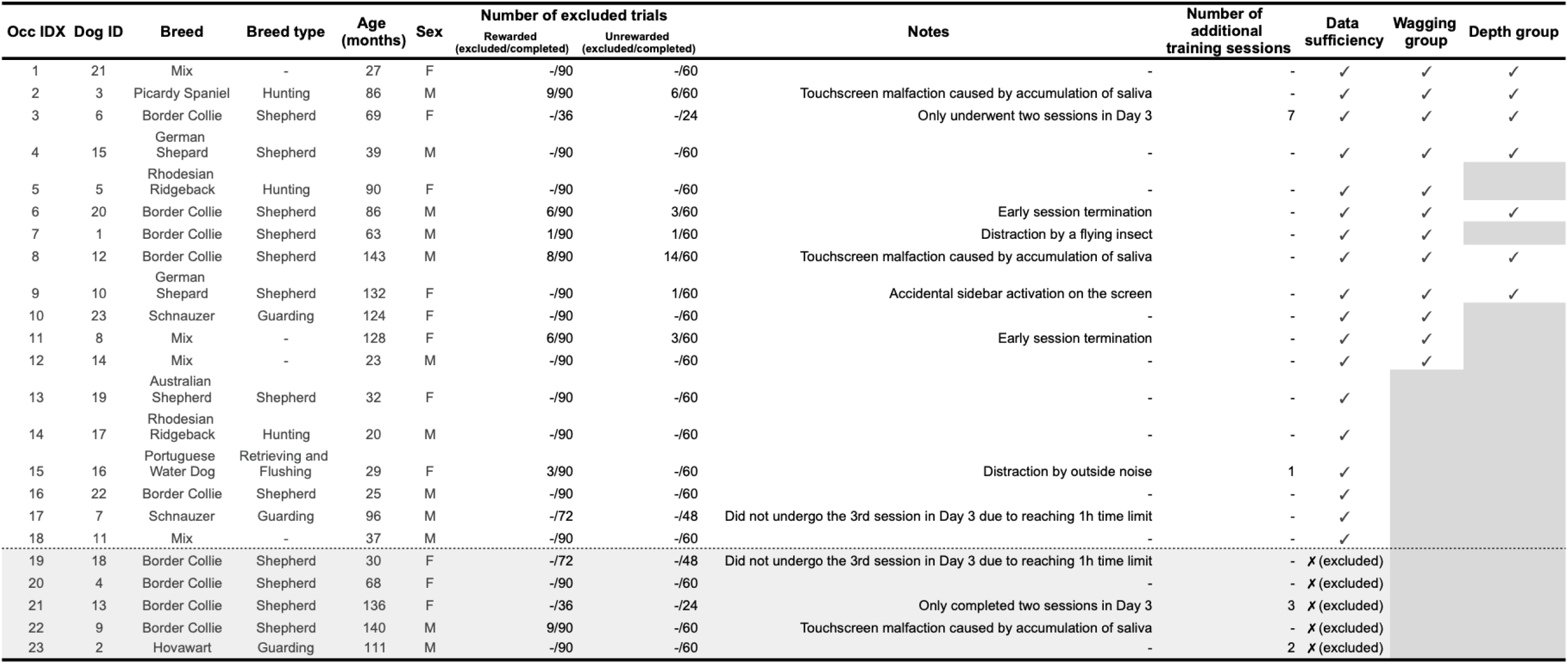

## METHOD DETAILS

### Ethic Statement

All procedures used in the present study were approved by the Ethics and Animal Welfare Committee of the University of Veterinary Medicine Vienna (protocol number ETK-022/02/2024). We obtained informed consent from all the dog owners prior to participation in this study.

### Experimental Setup

The experiment was controlled automatically using a touchscreen-based custom made experimental apparatus (**Figure 1A**, left) that included a 17’ computer monitor, an infra-red touch frame and a custom automatic feeder (Ajuwon et al., 2025). The experimental room was divided into two areas: a testing area (2.40 m × 3.39 m) and a waiting area (**Figure 1A**, right). The testing area was aligned and centred directly below a ceiling-mounted camera. The back side of the apparatus was enclosed by two extendable baby fences (0.31 × 2.00 × 1.03 m; Mendler) positioned next to the touchscreen-based testing apparatus. This arrangement was devised so that the dog’s moving space was limited to increase the opportunity for successful interactions with the apparatus while keeping the video field of view restricted to the area where a dog and the apparatus were present (the owner was not recorded). Owners were seated in the waiting area to mitigate dogs’ potential stress and anxiety caused by separation, but were instructed not to interact with their dogs during testing sessions (i.e., no eye contact, petting or verbal encouragement). To further minimise unintentional cues, owners wore sunglasses and were asked not to make direct eye contact with their dogs. These measures reduced potential external influences unrelated to the experimental procedures, ensuring standardised testing *Conditions*.

### Task Control

All task contingencies were controlled using a custom-made Bonsai workflow (Lopes et al., 2015, 2021). This included all task events (e.g., stimulus presentation) but also saved all non-video data, including screen interactions.

### Experimental Procedure

For each dog, the entire experiment took place over three days – one or two days for training and one or two days for testing, depending on each dog’s training progress. On *Day 2*, two testing sessions were conducted if the training was completed on *Day 1*. On *Day 3*, two or three testing sessions were carried out (maximum 1hour per day/dog). A training session (20 *Rewarded* trials) preceded the first testing session of the first testing day. The intervals between *Day 1* and *Day 3* ranged from 2 to 35 days (mean: 14.3 days). Two dogs underwent only two testing sessions on *Day 3* due to additional training, another two dogs only completed four sessions within the 1-hour daily time limitation, and the remaining 19 dogs completed all five sessions (see Individual dog information).

### Training Protocol (Day 1)

Dogs were trained to use the touchscreen-based testing apparatus and to initiate trials through a two-step training process. Upon entering the experimental room, a dog was allowed to explore the environment freely for 3 minutes. During training, the owner was allowed to be in the testing area (**Figure 1A**, right) and to interact with their dog.

Each dog underwent a pre-training session in order to successfully start trials and trigger food rewards using a shaping procedure in combination with positive reinforcement, where every screen touch resulted in a food reward. At the start of each trial, a black plus symbol appeared on a white background (*Available*). The first touch after trial availability triggered auditory feedback (a click sound) and the display of a blue or yellow circle (counterbalanced across animals) on a white background (*Reward Signal*). Following this, a reward was delivered instantly from the food exit at the bottom of the testing apparatus (Figure 1A, left) and was accompanied by a bell sound (secondary reinforcement), after which, the screen would once again be *Available*. If dogs failed to interact with the screen on their own, the owner and/or experimenter would enter the testing area to lure the dogs towards the touchscreen and food delivery was controlled manually by the experimenter in order to only reward when dogs touched the screen deliberately (i.e., not accidentally). This procedure (*Available* – *Touch* – *Reward Signal* – *Reward*) was repeated for 20 successful (without human intervention) trials.

The training sessions followed the pre-training. Every trial started with the screen turned to white with a centrally placed black plus symbol – the *Available* phase. When the dog touched the screen during the *Available* phase, the *Reward Signal* (blue/yellow) was presented for 1 second, marking the *Anticipation* phase. A food reward was then delivered with the bell sound, followed by the inter-trial-interval (*ITI)* where a black background lasting 3.5 seconds was presented and during which the screen was unresponsive to touches, yielding no consequences (**Figure 1B**). If a dog did not touch the *Available* screen within 20 seconds, a trial was repeated, starting with a shorter *ITI* (1.5 seconds) up to a maximum of two times (correction procedure). Except during the *ITI*, any touches to the screen were followed by auditory feedback (a click sound). This procedure (*Available – Touch – Reward Signal – Reward – ITI*) was repeated for at least 2 sessions with 20 trials each (excluding correction trials, see above) for each dog and was continued until the dog reliably touched the *Available* screen with their noses (or paws). A 5-minute break separated these sessions, during which the dog and owner waited outside the experimental room. Additional training sessions/days were added if dogs did not reach the criterion on *Day 1* (i.e., voluntarily and consecutively initiating trials without any guidance/assistance from the owner or the experimenter, see **Individual dog information** for details). All dogs underwent one training session prior to the first testing session of the first testing day.

### Testing Protocol (Day 2-3)

We adopted two experimental *Conditions* in testing sessions: the *Rewarded Condition*, where dogs could receive food following a 2-second waiting period, and the *Unrewarded Condition*, where no food reward was delivered after the same delay (Figure 1C, bottom). The testing protocol (Figure 1B) was similar to the training protocol but with *Anticipation* time increased from 1 to 2 seconds and the inclusion of the *Unrewarded Condition*. Based on a pilot experiment (data not shown), the *Anticipation* time was set to 2 seconds to prevent animals from disengaging from the task and to minimise the risk of frustration caused by longer delays (Amsel, 1958).

The first testing session on each day was preceded by 5 rewarded ‘warm-up’ trials. *Rewarded* and *Unrewarded* blocks alternated every 6 trials, for a total of 30 trials, with each session starting and ending with a *Rewarded* block. To signal the transition from *Unrewarded* to *Rewarded* blocks, a free reward (but no bell sound) was delivered 2.5 seconds after the last *Unrewarded* trial regardless of any responses to the screen. Sessions were terminated if dogs stopped responding to the task or consuming rewards for more than 3 consecutive trials in a *Rewarded* block. Specifically, the first session in *Day 1* was terminated for two dogs, from the 22^nd^ trial onward (see Individual dog information).

In total, there were 150 planned experimental trials (30 trials × 5 sessions) for each dog, excluding the daily warm-up phase. This included 90 *Rewarded* trials (6 trials × 3 blocks × 5 sessions) and 60 *Unrewarded* trials (6 trials × 2 blocks × 5 sessions). As in the training sessions, a 5-minute break separated each session, during which dogs and their owners were asked to leave the experimental room.

### Data Acquisition

We collected trial start times directly from the *Bonsai* program (Lopes et al., 2015) running the touchscreen-based testing apparatus (**Figure 1A**, left). Additionally, we recorded RGB-D videos at a resolution of 1280 × 720 pixels at 30 frames per second (fps) with a single depth camera (Azure Kinect DK, Microsoft) mounted on the ceiling (**Figure 1A**, right). Video data was captured via the Azure Kinect DK recorder software. For each trial, video data was divided into two distinct, non-overlapping task *Epochs*: 1) *Pre-Response*: 2 seconds before initiating a trial and 2) *Anticipation*: 2 seconds after initiating a trial (**Figure 1C**, top). These two *Epochs* were defined to investigate tail movements associated with anticipation states (*Prediction 2*) and also to ensure data consistency and comparability across *Conditions* and dog participants. Specifically, we did not include periods right after the outcome was revealed due to a high probability of ongoing food consumption during the *Rewarded Condition* interfering with tail movement readouts. Additionally, to account for variability in trial start times, we aligned the task *Epochs* to a dog-touch to the screen, not the start of each trial. Each task *Epoch* was identified based on RGB values in the stimulus presentation area of an upward-facing laptop screen that mirrored the content shown on the screen facing the dog and was visible to the ceiling-mounted camera.

### Trial Start Times

The trial start time was defined as the duration from the onset of *Available* to the moment a dog touched the screen (Figure 1B). If a dog did not touch the screen within the 2-trial correction procedure, the trial start time was determined as the maximum trial duration, 63 seconds (20 seconds for *Available* × 3 trials + 1.5 seconds for shorter ITI × 2).

### Pose Tracking from RGB Videos and Data Processing

The RGB videos were used to track the position of 7 points along the dog’s body using DeepLabCut 2.3.10 (Mathis et al., 2018). These points included nose, head, withers, back (centre of body), tail base, tail middle, and tail tip (**Figure 2A**). The last four points (back, tail base, tail middle and tail tip) were used for further analyses. We trained pose-tracking models for each dog to accommodate variations in size, colour, and body shape across the recruited dogs. Between 20 and 125 frames were randomly (or manually) extracted from the videos for initial labelling. Each model was trained for up to 100,000 iterations per training sequence using ResNet-50 (default). We analysed the videos and obtained each body’s XY coordinates across all frames (inference procedure). After every training, the results were visually inspected. Additional frames were labelled and followed by additional training until it was visually confirmed that most body points were accurately and consistently tracked when they were visible across all the frames; the mean of test error with p-cutoff (0.80) across all dogs’ final tracking models was 5.53 px (∼20mm) in the real environment. Each body point’s XY coordinates were exported and coordinates with low likelihood scores (< 0.80) were removed. Subsequently, all tracking data from DeepLabCut were processed using custom Python code (code will be made available upon acceptance). Firstly, frame-by-frame differences for each body point were calculated and outliers (i.e., discontinuities) were detected using an exponentially weighted moving average (EWMA) with a 120-frame window and a 3×SD (Standard Deviation) threshold, and replaced by NaN values. Then, up to 20 consecutive empty frames were linearly interpolated and the resulting data were smoothed using a Savitzky-Golay filter. The missing value rates for each tracked body point after the data processing described above (i.e. handling outliers, linear interpolation, and smoothing) can be found in Supplementary Table 1.

### Depth Data Alignment and Processing

#### a) Depth coordinate alignment

Depth images were acquired through the Azure Kinect DK with the NFOV 2 × 2 Binned mode (320 × 288 pixels) at 30 fps, the same frame rate as the colour images. Depth data were transformed to compensate for the 6-degree downward angle relative to the colour camera and to compensate for the different image resolutions. Additionally, depth values were re-aligned to the gravity coordinate system to reflect the actual distance from the camera to the experimental room’s floor (2380 mm, manually measured). Firstly, a depth frame without a dog was obtained and transformed into the colour coordinate system using the officially provided transformation function from Azure Kinect SDK (*k4a_transformation_depth_image_to_color_camera()*). The plane in the image was estimated using the RANSAC algorithm (Fischler and Bolles, 1981), resulting in a plane equation for the colour coordinate system, which served as a proxy for the room’s floor. To ensure the floor height matched the target value of 2380 mm, a projection matrix was derived so that points on the estimated plane corresponded to 2380 mm. This projection matrix was then applied to all data frames to correct the depth values (see Data processing diagram).

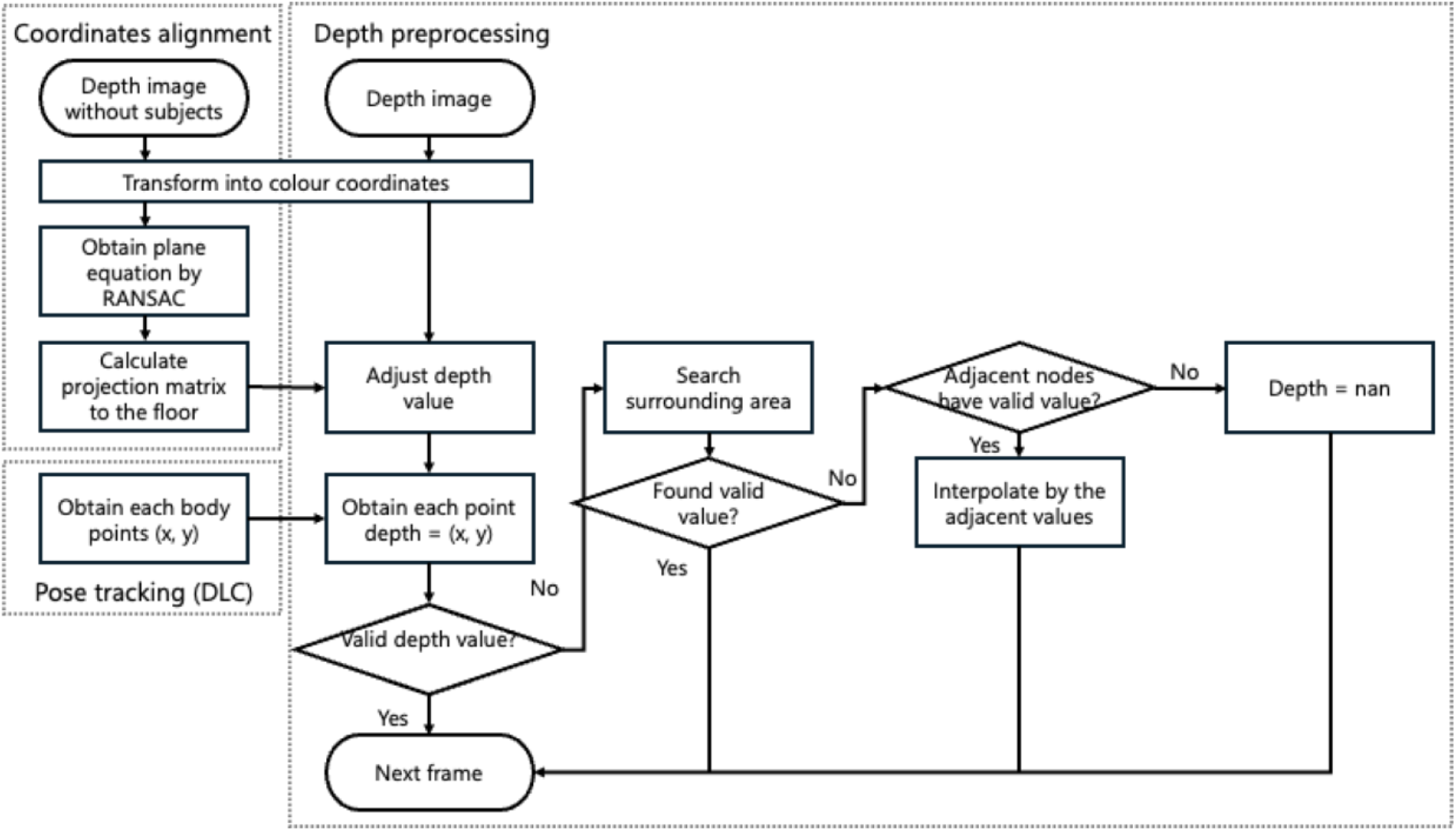

### Data processing diagram

The depth data (320 × 288 pixel resolution) were first transformed into the colour coordinate system (1280 × 720 pixel resolution), followed by the application of a projection matrix to align the coordinate system parallel to the ground floor plane (*Coordinates alignment*). Subsequently, the depth values corresponding to each tracked body point were extracted. If a depth value on the tracked point’s XY coordinates was invalid, a valid value identified by searching the surrounding area of the invalid point was adopted or a depth value was interpolated by adjacent nodes’ depth values (*Depth preprocessing*).

#### b) Depth data processing

We obtained the processed depth value for each body point by accessing the X and Y coordinates of the tracked points in the colour frame. However, the Azure Kinect DK measures the depth indirectly using the Time-of-Flight (ToF) principle; hence, the depth values of the tracked body positions may occasionally be invalid. To address this issue, we searched the surrounding area of invalid points with a radius of up to 7pixels (∼25 mm in the real environment) and adopted the highest depth value from the ground level. If no valid value surrounding the invalid point was found, but if adjacent body points had valid depth values, the invalid value was linearly interpolated using the adjacent nodes’ values. Similar to the processing of tracking data, outliers in the depth time series data were detected using the EWMA method (frame window: 120 frames, threshold: 2.5 × SD) and replaced by NaN. Unlike the pose-tracking data, frame-by-frame differences were not used to detect outliers in this case, as the original depth data contained missing values more frequently than the tail angle data, which might have resulted in the false detection of non-outliers. Missing values were interpolated and all data were smoothed using the same procedure as for pose-tracking data. Finally, the depth values from the camera were converted to heights relative to the ground level by subtracting the original depth values from 2380 mm. Refer to Supplementary Table 1 for the missing value rates of each body point following data processing.

### Tail Kinematic Parameters

We calculated six parameters to describe tail movement kinematics: angle, wagging rhythmicity, wagging amplitude, wagging velocity, wagging acceleration, and height (**Figure 2B**). As a proxy for tail angle relative to the dog’s body, we used the angle between two vectors: (1) the vector defined by back and tail base points and (2) the vector defined by the tail base and the tail middle points. Tail middle was used instead of tail tip as we considered the tail middle to be more robust than tail tip (the vector defined by the tail middle is a better approximation of the overall tail position than the one defined by the tail tip, which can be heavily affected by slight deviations of the tip’s position and also less reliably tracked (tail middle also had a lower percentage of missing values; see Supplementary Table 1). Negative to positive values indicate left to right positions (**Figure 2A**). Tail-wagging rhythmicity was quantified using the entropy of the normalised power spectrum of the tail angle time series within a 2-second analysis task *Epoch* following a Fourier transform. Smaller entropy values indicate more rhythmic waves (i.e., composed of more consistent frequency bands). Tail wagging (peak-to-peak) amplitude was defined as the difference between adjacent positive and negative peaks, identified using Scipy’s *find_peak()* function. Tail wagging velocity was calculated by dividing frame-by-frame differences in tail angle by 1/30 seconds, and tail wagging acceleration was derived from velocity similarly. Tail height was defined as the height of the tail middle point relative to the floor (see Section *Depth Data Alignment and Processing* for details). Statistical analyses of rhythmicity, wagging velocity, and acceleration were conducted only for trials in which tail wagging was detected (see *Data Selection - Quantification of tail wagging occurrence* for more details on tail detection). All parameters were obtained separately for each task *Epoch* (*Pre-Response/Anticipation*).

## QUANTIFICATION AND STATISTICAL ANALYSIS

### Data Selection

#### Data exclusion

The first trial in each block was excluded to mitigate for potential heighted responses following block transitions. Additionally, some trials were also excluded from analyses due to early termination of sessions, touchscreen malfunctions caused by dogs’ saliva, and distraction caused by ambient noise. The proportion of excluded trials across all dogs was 2.18% for both Rewarded (42/1926 trials), and Unrewarded (28/1284 trials) *Conditions* (See Individual dog information for details). For tail height, we excluded data points when the tail base height was below a threshold calculated individually for each dog using Otsu’s method (i.e., that determines the optimal threshold by maximising the between-class variance, and at the same time minimising the within-class variance; for details see Otsu, 1979) to filter out non-standing postures (e.g., sitting or crouching). This resulted in an average 14.2% datapoints excluded (*min* = 2.2%; *max* = 34%) and allowed for a consistent comparison of tail height in standing dogs across all *Conditions* and task *Epochs*.

#### Tracking quality evaluation

Before analysing tail kinematic parameters, we evaluated the tracking quality based on missing value rates. First, each dog’s missing value rate for each body point was calculated for each task *Epoch.* Specifically, this was defined as the ratio between the total number of missing frames for a given body point, and the total number of frames within the corresponding epoch of all the valid trials. Then, the average missing value rate for each body point was obtained across the two task *Epochs* (Supplementary table 1). Dogs with an average missing value rate above 50% for the tail middle, tail base, or back (i.e., the points used for tail angle calculation) were excluded from the tail kinematic analysis (*n =* 5; Occ IDX: 19-23; see Individual dog information). Visual inspection of the video of these dogs confirmed minimal to no tail movement during the experiment. The 18 remaining dogs were included in the tail kinematics analysis.

#### Quantification of tail wagging occurrence

We first quantified tail-wagging occurrence during testing sessions. Here, we defined tail wagging as tail angle movements with a peak-to-peak amplitude of at least 20 degrees and a frequency of at least 1 Hz. A sinusoidal wave, with a peak-to-peak amplitude of 20 degrees and a frequency of 1 Hz, sampled at 30 fps, was generated for 2 seconds (60 frames). Its Fourier transform revealed a power of approximately 2900 in the 1 Hz frequency band. Based on this result, we established the following criteria to determine the presence of tail wagging within a 2-second analysis *Epoch*: (1) the power in the 1Hz band ≥ 2900, or (2) a peak in the frequency range above 1 Hz detected by the Scipy’s *find_peak()* function with power ≥ 2900. Tail-wagging occurrence ratios, defined as the proportion of trials in which tail-wagging occurred out of valid trials across 2 task *Epochs* (*Pre-Response and Anticipation*), were calculated for each dog and ranked in descending order. The minimal tail wagging occurrence threshold was set at 0.25, with dogs with occurrence ratios above it categorised as the wagging group (**Figure 3A**).

#### Curated datasets

Tail parameter analyses were carried out only on the wagging group dogs (6 males and 6 females; mean age: 7 years (84 months), range: 23 – 143 months). For the tail height analysis, 7 dogs (4 males and 3 females; mean age: 7 years (83 months), range: 27 – 143 months) were used from the wagging group. Three dogs were excluded due to infrared absorption caused by black fur, and the other two due to their tails being too thin to allow for reliable capture of depth values.

#### Statistical Analyses (models)

All analyses were conducted using RStudio (version 3.6.0+; R Core Team, 2018), and results were considered significant for p < 0.05. A series of linear mixed-effects models (LMMs) were employed using the “lmer” function from the “lme4” package (Bates et al., 2003) for the continuous response variables: latency to start the trial (i.e., trial start times) and tail kinematic parameters. For binary outcome variables: the occurrence of tail wagging (Binary response values; 0 or 1), generalized linear mixed models (GLMMs) with a logit link function were fitted using the “glmer” function.

Separate models were fitted on two sets of measures: the latency to start a trial (*Trial Start Times*) and the different tail kinematic parameters of interest, which included wagging occurrence, angle, rhythmicity, amplitude, velocity, acceleration, and height. Prior to modeling, the residuals of all variables were visually inspected using histograms and QQ-plots to check distributional assumptions and identify missing values.

For each measure, the effects of interest were the task *Condition* (*Rewarded vs. Unrewarded*), the *Epoch* (*Pre-Response vs. Anticipation*) except for *Trial Start Times*, the *Session* number and the *Trial* progression, which were included as predictors in the models.

To account for individual variability we included random intercepts and random slopes for these predictors across dogs (*Dog ID*).

To keep the 5% type I error rate when testing the overall effect of the predictor variables on the response variable, we compared the “full” models, which include the predictor variables of interest, with a simpler “null” model that excludes them. These comparisons were performed using likelihood ratio tests (function “anova”; Dobson, 2001).

All the covariates were centered, or z-transformed, to enhance model convergence and interpretability. The model diagnostics involved assessing residual distributions and the normality of the “Best Linear Unbiased Predictors” (BLUPs: “ranef” function, package “lme4”) – the model-derived individual-specific estimates of random effects. These represent deviations of each dog from the population average, ensuring that the assumption of normally distributed random effects was confirmed, thus supporting the validity of the model. Furthermore, model stability was evaluated by checking for multicollinearity through Variance Inflation Factors (VIF – “vif” function, library “car”) and all values remained well below the common threshold (VIF < 2).

Model stability was evaluated iteratively by excluding one level of the random effect (i.e., one dog) and then refitting the model to check whether the fixed effect estimates changed substantially (glmm.model.stab; Nieuwenhuis et al., 2012). No significant fluctuations were observed, implying the stability of the model.

Final models were refitted using restricted maximum likelihood (REML), which helped provide more accurate estimates of the model’s effects. When needed, post-hoc pairwise analyses were performed using estimated marginal means (“emmeans” function from the “emmeans” package; Lenth and Piaskowski, 2017) to allow for a detailed exploration of the levels of the factors in the models: *Condition*, *Epoch* and, when relevant, their interaction.

